# HySA: A Hybrid Structural variant Assembly approach using next generation and single-molecule sequencing technologies

**DOI:** 10.1101/069815

**Authors:** Xian Fan, Mark Chaisson, Luay Nakhleh, Ken Chen

**Affiliations:** Department of Computer Science, Rice University, Houston, Texas, USA; Department of Bioinformatics and Computational Biology, Division of Quantitative Sciences, The University of Texas MD Anderson Cancer Center, Houston, Texas, USA; Department of Genome Sciences, University of Washington School of Medicine, Seattle, Washington, USA

## Abstract

Achieving complete, accurate and cost-effective assembly of human genome is of great importance for realizing the promises of precision medicine. The abundance of repeats and genetic variations in human genome and the limitations of existing sequencing technologies call for the development of novel assembly methods that could leverage the complementary strengths of multiple technologies.

We propose a Hybrid Structural variant Assembly (HySA) approach that integrates sequencing reads from next generation sequencing (NGS) and single-molecule sequencing (SMS) technologies to accurately assemble and detect structural variations (SV) in human genome. By identifying homologous SV-containing reads from different technologies through a bipartite-graph-based clustering algorithm, our approach turns a whole genome assembly problem into a set of independent SV assembly problems, each of which can be effectively solved to enhance assembly of structurally altered regions in human genome.

In testing our approach using data generated from a haploid hydatidiform mole genome (CHM1) and a diploid human genome (NA12878), we found that our approach substantially improved the detection of many types of SVs, particularly novel large insertions, small INDELs (10-50bp) and short tandem repeat expansions and contractions over existing approaches with a low false discovery rate. Our work highlights the strengths and limitations of current approaches and provides an effective solution for extending the power of existing sequencing technologies for SV discovery.

## Introduction

Completely, accurately and cost-effectively assembling human genomes is a prerequisite for genomic medicine. Advances in translational genomics are hampered by technical challenges in assembling structurally altered regions in human genome, which are shown to be essential for generating genetic diversities and human diseases (Feuk et al. 2006; Sharp et al. 2006; Lupski 2007). Advances in Next Generation Sequencing (NGS) technologies have greatly facilitated assembly and detection of structural variations (SV) in human genome (Alkan et al. 2011a). Many computational methods have been developed to identify SVs through examining alignments of paired-end reads to the human reference genome, scanning for abnormally aligned reads (such as unmapped reads, discordant read pairs, clipped reads and reads with large gaps) and variation of read depth, and inferring SV positions and orientations (Chen et al. 2009; Wang et al. 2011; Layer et al. 2012; Rausch et al. 2012; Sindi et al. 2012). Others perform whole-genome or targeted assembly of sequencing reads and identify SVs from pair-wise alignment of assembled contigs against the reference (Iqbal et al. 2012; Chen et al. 2014; Xie et al. 2014). However, although NGS reads have low base-calling error rates (Ross et al. 2013), their read lengths are often limited (e.g 100-200 bp for Illumina Hi-seq instruments). The short read length leads to a bias against assembly and detection of SVs, which often occurs near segmental duplications or large repeats in the genome (Alkan et al. 2011b). Moreover, complex sequence alterations around SV breakpoints (e.g microhomology and micro-indels (Hackl et al. 2014), Kataegis (Alexandrov et al. 2013), etc.) substantially hamper sensitivity and specificity of SV detection methods that depend on aligning individual reads against the reference (Consortium 2010).

The advent of Single Molecule Sequencing (SMS) technologies greatly changed the landscape of genome assembly approaches as they provide much longer reads (e.g., 12kb on average for Pacbio reads in P6-C4 chemistry). As a result, many SVs missed by NGS can now be detected (Chaisson et al. 2015b). Unfortunately, the SMS technologies are typically error prone due to the use of one molecule for real-time sequencing. For example, Pacbio reads have an error rate at 15%, and the majority (14%) of the errors are indels (Ross et al. 2013). The high error rate has posed new challenges to bioinformatics tools that perform alignment or assembly. To tackle the challenges, BLASR (Chaisson and Tesler 2012) was developed to tolerate the INDEL errors in aligning error-prone Pacbio reads to high quality sequences in a computationally efficient way. In addition, the widely used BWA algorithm (Li and Durbin 2009) was also extended to align Pacbio reads (Li 2013). It is plausible to infer SVs based on an analysis of long read alignment to the reference by searching for INDEL signals (gaps inside a read alignment) and stop signals (clipped ends) (English et al. 2014; Chaisson et al. 2015b). However, it is often difficult to accurately assign gaps or stops due to high error rate, long read length and prevalence of repeats. As a result, it is challenging to parse Pacbio read alignment to accurately determine SVs and breakpoints, particularly those resulting from novel (non-reference) insertions.

On the other hand, *de novo* genome assembly approaches have rapidly advanced in terms of both quality and computational efficiency (Chaisson et al. 2015a). Overall, three different paradigms have been developed 1) overlap-layout-consensus (OLC) (e.g., Mira, Newbler, Celera Assembler (Myers et al. 2000)), 2) *De Bruijn* graph (e.g., Velvet, SOAPdenovo, AbySS, and ALLPATHS) and 3) string graph (Myers 2005). The *De Bruijn* graph based approaches require high quality reads and are only applicable to NGS data, while the OLC and string graph based approaches are applicable to both NGS and SMS data. For example, Falcon (Chin, C.-S., P. Peluso, F. J. Sedlazeck, M. Nattestad, G. T. Concepcion, A. Clum, C. Dunn, R. O′Malley, R. Figueroa-Balderas, A. Morales-Cruz et al. 2016), a recently developed assembly algorithm utilizes string graph to assemble a diploid genome from Pacbio reads. Although the construction of string graphs takes a linear time using FM-index, Falcon is computationally intensive due to an error correction step that requires pairwise alignment of all Pacbio reads. Chin et al. (Chin et al. 2013) also utilized relatively short Pacbio reads to correct long Pacbio reads to facilitate assembly. Berlin et al. (Berlin et al. 2015) made use of a statistical hashing technique for pairwise overlapping, which greatly reduces computation time. For the purpose of SV detection, however, *de novo* whole genome assembly (WGA) is not optimal since the majority of the genome does not contain SVs. They often require computational resources not widely available when assembling large mammalian genomes (∼3 Gbp) (Pendleton et al. 2015). Moreover, they are often not optimized to assemble diploid genomes containing heterogeneous SVs. In comparison, targeted SV assembly approaches (Chen et al. 2014) that aim to assemble sequences spanning SVs are often more effective in terms of computational efficiency and SV detection power, as it dissects a WGA problem into a set of independent local assembly problems that can be more effectively solved. However, in addition to performing powerful local assembly, targeted approaches need to 1) achieve comprehensive unbiased selection of targets and 2) ensure the results obtained from local solutions are also globally optimal.

Considering the advantages and disadvantages of the technologies, i.e., NGS reads are short but accurate, whereas SMS reads are long but inaccurate, a hybrid assembly approach that combines data from the two or more technologies can potentially achieve more powerful assembly and SV detection. Ideally, the accuracy of NGS reads can be used to correct errors in SMS reads, whereas the length of SMS reads can be used to confidently anchor the assemblies to the reference. A few efforts aiming to achieve such combination, although not specifically for SV detection, have been proposed. A toolbox has been developed to simulate the integration of multiple technologies for optimal personal genome assembly (Du et al. 2009). PacbioToCA (Koren et al. 2012) performs hybrid *de novo* WGA by aligning all NGS short reads to all Pacbio reads for error correction. LSC (Au et al. 2012) applies a similar strategy but aimed to reduce the error rate in homopolymer runs. While these methods utilize the high fidelity of NGS reads and the long length of Pacbio reads, they turn to be computationally intensive and are not designed for SV detection. MultiBreak-SV (Ritz et al. 2014), on the other hand, uses a probabilistic approach that can combine the alignment of individual SMS and NGS reads for detecting SVs, particularly regions involving multiple SVs. However, it is not suitable to detect novel insertions since it relies on accurate alignment of individual reads to the reference. Moreover, it only examines discordant aligned NGS read pairs (but not unmapped or clipped NGS reads).

In all, there lacks a computationally efficient hybrid assembly approach for accurate SV detection despite a pressing demand in applied genomics. The main obstacle is a lack of computationally efficient algorithms that can effectively synergize heterogeneous data sources of highly discrepant properties (e.g., read length and sequencing error) with highly structured contents (e.g., sequence homology in human genome)

In this manuscript, we propose a novel Hybrid Structural variant Assembly (HySA) method that identifies and performs genome-wide SV assembly from both NGS and SMS data. Our method targets genomic regions that cannot be accurately mapped by short reads and utilizes long reads to resolve structural complexities. It alleviates the challenges in assigning gaps in alignments of error-prone long reads and in aligning long reads containing non-reference sequences. We show using both simulated and real data that our method can identify an appreciate amount of novel SVs and can effectively complement existing approaches for SV detection.

## Results

### Method overview

Our HySA method (**Fig. 1a**) requires two sets of input data: Set A of reference alignment of paired-end short reads generated by low-error-rate NGS (such as Illumina HiSeq) and set B of long reads generated by high-error-rate SMS (such as PacBio SMRT-seq). It first identifies and extracts unmapped, discordantly paired and end-clipped short reads in set A and then aligns them to the set of long reads in set B (**Methods**). The set of aligned short and long reads form a bipartite graph, in which one set of nodes represent short reads, the other represent long reads, and edges between them represent confident pair-wise alignments. The extracted short reads are often from disjoint regions of unique sequence context due to sparseness of SVs and short fragment size of set A.

**Fig 1.**
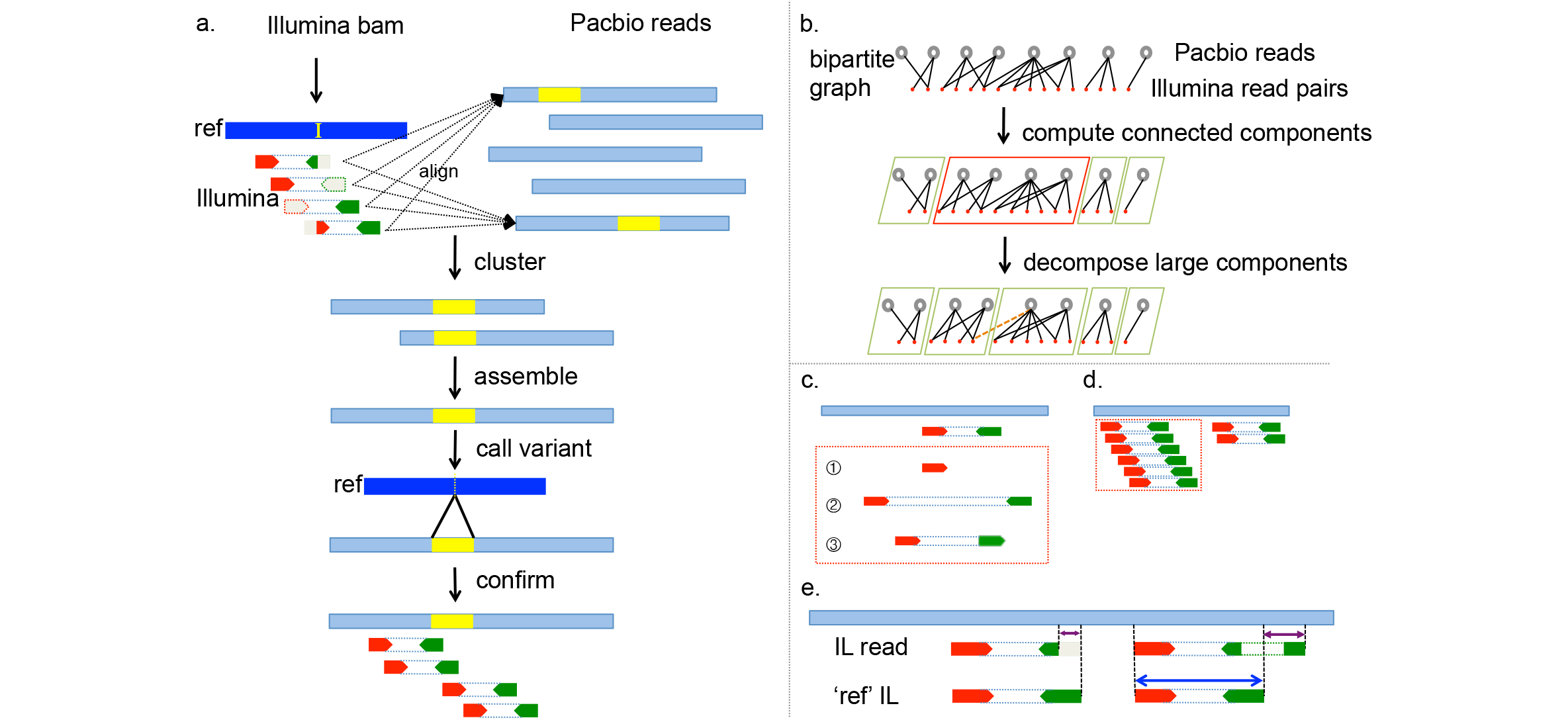
Diagram of the HySA approach for SV assembly and detection. **a).** Abnormally aligned Illumina reads are extracted from a BAM file and aligned to a set of Pacbio reads (light blue) generated from the same DNA sample. The cluster of reads associated with an SV is identified using a set of bipartite-graph partitioning algorithms. Contigs are assembled from Pacbio reads in each cluster and are aligned to the reference, from which SVs and breakpoints are identified and further confirmed by Illumina reads in the same cluster. An insertion (the yellow segment in reference and the yellow segments in Pacbio reads) is used for illustration. The Illumina reads are in red and green, corresponding to the forward and backward strands, respectively. The subsequence or the whole read that cannot be mapped is in grey. **b**) Clustering strategy. A bipartite graph is built from the pair-wise alignment of Illumina reads to Pacbio reads. One set of the nodes corresponds to Pacbio reads (top row, black open circles), and the other set corresponds to Illumina read pairs (second row, red solid circles). An edge is added when there is a reliable alignment between an Illumina read pair and a Pacbio read. The bipartite graph is decomposed into connected components (green and red boxes) using, e.g., a Union-Find algorithm (Sedgewick and Wayne 2011). Large components, e.g., the one in the red box, are further decomposed into communities of expected sizes using a graph decomposition algorithm called Infomap (Rosvall and Bergstrom 2008). **c**) False alignments between Illumina and Pacbio reads are illustrated in dashed red box: 1) single end alignment; 2) paired end with abnormal insert size; and 3) paired end with abnormal orientation. **d**) False alignments due to repeats. Illumina reads’ alignments against a Pacbio read (in dotted red box) are filtered out when the depth of Illumina reads significantly exceeds the expected coverage. **e**) A competitive alignment strategy that eliminates false alignments between Pacbio and Illumina reads. For each Illumina read pair, a pseudo (ref) read pair is synthesized from the reference sequence in identical positions and orientations. An alignment between an Illumina read pair and a Pacbio read is false when 1) the Illumina pair has a shorter aligned sequence against the Pacbio read than does its pseudo pair or 2) the alignment of the Illumina pair implicates an abnormal insert size whereas the pseudo pair does not.

Consequently, the bipartite graph is often sparse and can be computationally efficiently (near linear complexity w.r.t. the number of nodes) decomposed into connected components (CC) using a Union-Find algorithm. Each CC often corresponds to one SV sequence containing at least 1 breakpoints with its size (number of nodes and edges) proportional to the average physical coverage in both sets A and B. False alignments between short and long reads could lead to inaccurate nodes and edges in the graph and result in erroneously large connected components (ELCC). When that occurs, we further decompose ELCCs into small communities via a network flow based graph algorithm (Rosvall and Bergstrom 2008) (**Fig. 1b**). This algorithm iteratively merges and splits small communities in a random order until they have expected sizes and no better partitioning can be found. Each resulting CC or community contains a cluster of long and short reads that are expected to come from a single genomic origin. Assembling long reads in each cluster into contigs and aligning them to the reference enable discovery of SVs. To reduce false discovery, short reads in the same clusters of the long reads are aligned to the assembled contigs to confirm the identified SVs (**Fig. 1a**).

Noticeable features of our algorithm include:

1. The read-clustering approach via partitioning of a bipartite graph is reference-agnostic, which allows reads containing novel non-reference sequences to be clustered and assembled together and thus facilitates the assembly of non-reference insertions;

2. Only a subset of potentially variant-containing reads is analyzed, which leads to substantial saving in computational cost, as compared with whole-genome assembly approaches; and

3. No direct alignment of individual long reads to the reference is needed. This not only reduces computation, but more importantly, alleviates difficulties in assigning gaps or stops when aligning noisy long reads to the reference.

### Quality of the bipartite graph

The quality of our approach is governed by the accuracy of the bipartite graph that we construct from the data. To assess the quality of our approach, we performed a simulation experiment using the nucleotide sequence of chromosome 21. We randomly selected 310 positions on chromosome 21 separated by at least 60 kbp and at least 30 kbp from any gap. We simulated paired end Illumina reads spanning the selected positions at a coverage of 5×, 10× 15×, 20×, 25× and 30× and a read length of 100, 125 and 150 bps, respectively, and Pacbio reads covering the whole chromosome 21 at a coverage of 10×,30× and 60×, respectively (**Methods**). We then aligned each set of Illumina reads to each set of Pacbio reads using BWA mem with the “pbread” option and BLASR, respectively and compared clusters of reads identified using our algorithms against the ground truth, i.e., clusters of reads derived from the known positions of the synthetic reads. We measured the accuracy of clustering using JI90 (**Methods**), a metric ranging from 0 to 1 with 1 indicating perfect inference of the clusters. We found that JI90 increases generally as coverage and read length increase and that BLASR led to considerably higher JI90 than BWA when Pacbio coverage increased to over 10× (**Methods** and **Supplemental Fig. 1**). With over 15× Pacbio coverage, the JI90 obtained with BLASR increased to greater than 0.6 in all Illumina coverage and read lengths. It further increased to 0.8 when Pacbio coverage is over 25x, indicating that BLASR and our subsequence bipartite graph partitioning algorithms can effectively hybridize Illumina and Pacbio reads and generate a reasonably accurate representation of the targeted regions in the genome.

### SV detection in a haploid genome CHM1

The performance of our algorithm can be measured based on SV detection sensitivity and specificity, in comparison with other SV detection algorithms on the same data sets. We ran our algorithm on 50× Illumina and 46× Pacbio whole genome sequencing data generated from a hydatidiform mole haploid genome (CHM1) (Chaisson et al. 2015b). SVs in this genome have been well characterized in previous studies using approaches that analyze BLASR alignment of Pacbio reads to the reference (Chaisson et al. 2015b). Moreover, a high quality *de novo* whole genome assembly constructed from Pacbio reads (Berlin et al. 2015) and further confirmed by an independent high coverage (200×) Illumina whole genome assembly (Steinberg et al. 2014) was available as a reference to validate our results.

Our algorithm extracted 0.28% of the 50× Illumina reads and 6.8% of 46× Pacbio reads. In all, 130,058 (72,354 from Union-Find, 57,704 from Infomap decomposed from one ELCC) clusters were formed and 114,230 (71,092 from Union-Find, 43,138 from Infomap) successfully assembled into at least one contig, which led to the detection of 32,121 SVs including 3,007 large deletions (size > 50bp), 4,587 large insertions (size >50bp), 12,401 small deletion (size ≤ 50bp) and 12,126 small insertion (size ≤ 50bp) (**Supplemental Table S1**). The two main steps of HySA, alignment and assembly took a total of around 36,000 CPU hours on a high performance BL465c G7 blade with AMD 6174 processors and less than 12 GB memory per node. Both the CPU and the memory cost were at least an order of magnitude lower than what were required to perform a *de novo* whole genome assembly (Chin et al. 2016).

### Large deletions in CHM1

Among the 3,007 large deletions we called (referred below as HALD), 2,557 (85%) were directly validated by aligning the assembled SV contigs to the Berlin et al. assembly (**Methods**). A detailed look at the calls that cannot be validated by the Berlin et al. assembly provided an estimated false discovery rate (FDR) of 7.5% (**Methods**). For comparison, we generated a merged call-set (MGLD) containing 2,645 deletions discovered by Delly (Rausch et al. 2012) from the 50× Illumina data and by Chaisson et al. (Chaisson et al. 2015b) from the 46× Pacbio data (**Methods**). Thus, the differences between HALD with MGLD can reveal the unique difference of our approach relative to a naïve approach that does not perform hybrid assembly but merges call-sets independently derived from a single technology. In total, 1,961 (74.1% of MGLD, 65.2% of HALD) were shared between these two sets (requiring 50% reciprocal overlap). Importantly, 659 (21.9% of HALD) deletions were uniquely discovered by our approach and were further validated using the Berlin et al. assembly, indicating a shear gain in discovery power attributable to our methodology.

We further found that these 659 deletions uniquely discovered by our approach are associated with significantly less variant-supporting Illumina reads (7.28 versus 29.28, P-value= 1.382e-14, Students’ T-test) than are deletions in the MGLD. Although the numbers of variant-supporting Pacbio reads (**Methods**) do not differ significantly between the two sets, the distributions of gap starting positions relative to the breakpoints were significantly different (mean: 18.06 bp versus 43.77 bp, standard deviations: 19.56 versus 42.24 bp, P-value < 2.2e-16, two-sample Kolmogorov-Smirnov test). This observation confirmed the challenges in accurately aligning Pacbio reads to the reference. It is difficult to obtain consistent gap opening positions and sizes from BLASR alignment due likely to the high error rates of Pacbio reads and repetitive sequence context. Overall, these statistics confirmed that the novel deletions discovered by our approach were indeed associated with weak signals in either Illumina or Pacbio data and thus difficult to be called by Delly or approaches in Chaisson et al. (Chaisson et al. 2015b)

**Table 1.**
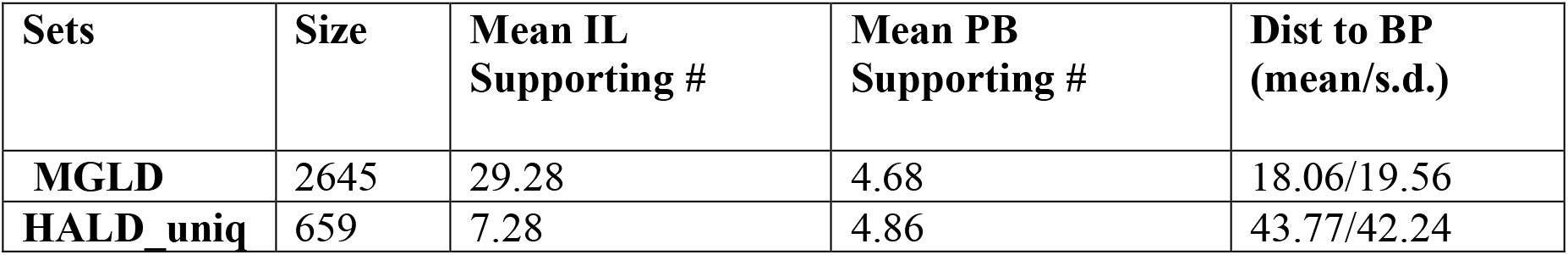
Comparison of calls from single technologies and those from the HySA approach. The two sets of calls (MGLD: merged calls from Delly and Chaisson et al.; HALD_uniq: calls unique to our hybrid method) validated by the Berlin et al. assembly are compared in terms of 1) mean Illumina supporting number, 2) mean Pacbio supporting number and 3) distance to the reference breakpoints (mean/standard deviation).

| Sets | Size | Mean IL Supporting # | Mean PB Supporting # | Dist to BP (mean/s.d.) |
| --- | --- | --- | --- | --- |
| **MGLD** | 2645 | 29.28 | 4.68 | 18.06/19.56 |
| **HALD_uniq** | 659 | 7.28 | 4.86 | 43.77/42.24 |

On the other hand, there were 684 deletions in MGLD missed by our approach due potentially to challenges in extracting and clustering the variant supporting reads.

Nonetheless, the results indicate that our approach can complement existing approaches and improve the overall discovery power.

### Large insertions in CHM1

Among the insertions (size > 50bp) detected by our method, 1,165 contigs contained inserted sequence longer than 500bp. Among them, 778 could not be aligned to the NCBI build v37 reference assembly and were novel (non-reference) insertions (**Methods**). Two hundred and eleven (211) of them could be aligned to the NCBI build v38 assembly, including 9 uniquely identified by our approach that were not reported by Chaisson et al. (Chaisson et al. 2015b). Among the 567 insertions that could not be aligned to the build v38, 522 could be aligned to the Berlin et al. assembly, including 20 uniquely identified by our approach that were not reported by Chaisson et al. Thirty-five of the rest 45 insertions that can be neither aligned to the Berlin et al. assembly nor aligned to build v38 were also reported by Chaisson et al., indicating their potential validity and the possibility of further improving the Berlin et al. assembly as well as the build v38 assembly using our results. Only 10 (1.3%) of the novel insertions discovered had no evidence of support from the available data.

In summary, our approach discovered 29 validated large novel insertions that were missed by Chaisson et al. This can be largely credited to our approach, which does not rely on having accurate alignment of Pacbio reads to the reference. For certain classes of insertions, it is easier to detect them from the alignment of Illumina read to the reference than from the alignment of long Pacbio read due to the accuracy of Illumina reads. Apparently, hybridizing Illumina and Pacbio reads together through our HySA algorithm has improved the assembly of insertions over approaches that involve error-prone reference alignments. An example of a 3 kbp novel insertion validated by NCBI build v38 is shown in **Supplemental Fig. 2**. As shown, Pacbio reads containing novel insertions could not be accurately aligned to the reference. However, they could be correctly clustered together through short Illumina reads (**Supplemental Fig. 3**), a large portion of which were both-end unmapped reads (blue vertical bars in **Supplemental Fig. 2b**) and one-end unmapped reads that can be partially anchored on the reference genome (red vertical bars in **Supplemental Fig. 2b**).

**Fig. 2.**
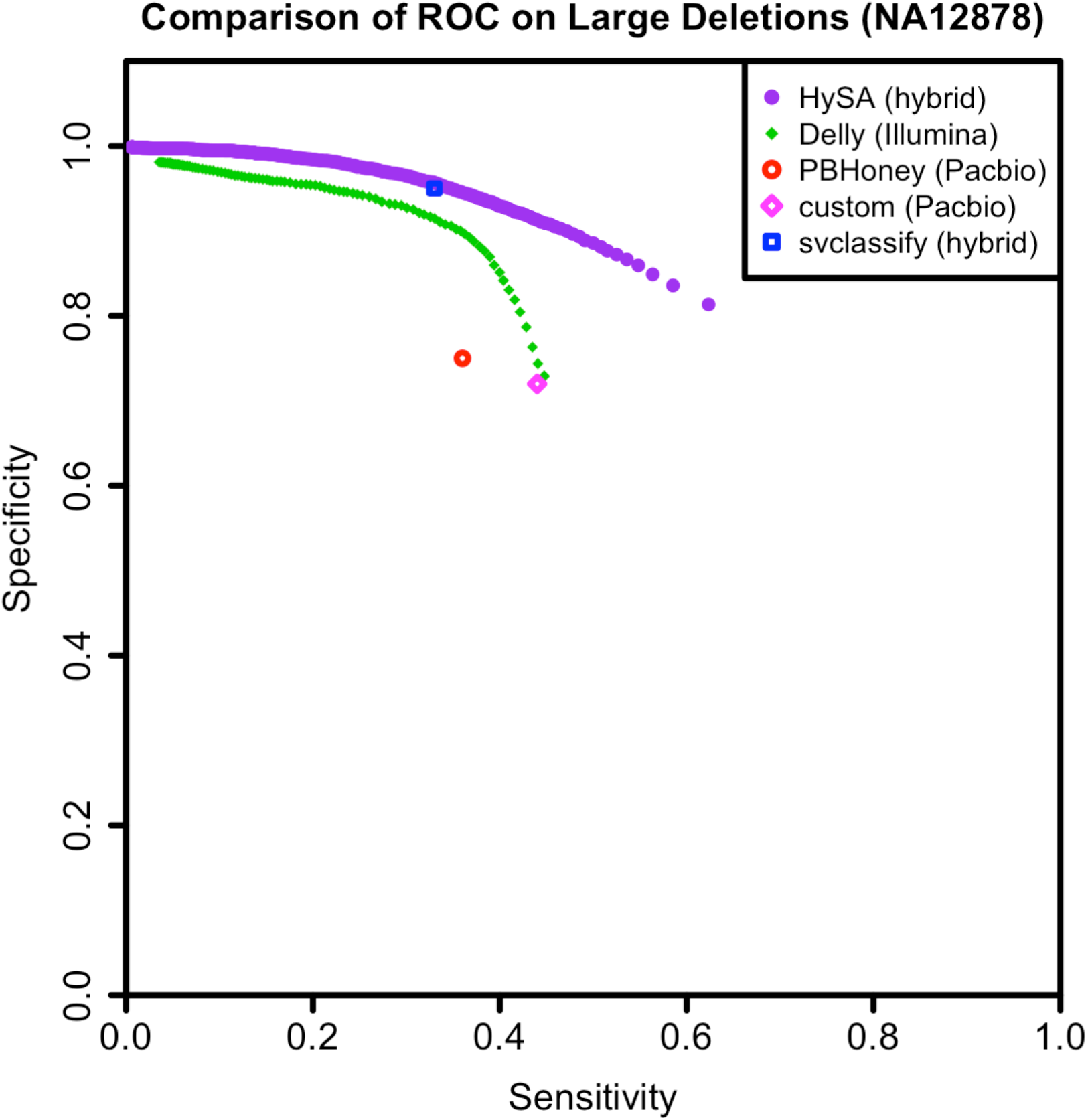
A comparison of detection sensitivity and specificity of large deletions in NA12878, among five competing approaches 1) HySA, 2) Delly, 3) PBHoney, 4) a custom pipeline based on Pacbio data alone and 5) svclassify.

### Small INDELs in CHM1

We compared small INDELs (≤ 50 bp) detected by our approach with those detected by Pindel (Ye et al. 2009) and GATK (DePristo et al. 2011) (**Methods**, **Supplementary Fig. 4**). The minimum size of the INDELs we detected was 11bp. A majority (95.1% for deletions and 84.5% for insertions) of our calls overlapped with either Pindel or GATK, and a large portion (85.9% for deletions and 68.4% for insertions) overlapped with both, showing the specificity of our call set. Notably, our approach was able to identify 2,538 novel INDELs that were missed by Pindel or GATK. The majority of these novel INDELs (92% deletions and 90% insertions) overlapped with the known INDELs in the dbSNP (Sherry et al. 2001) or with locations and sequence motifs indicating expansions or contractions of short tandem repeats (STRs) in the reference.

### SV detection in a diploid genome NA12878

We further examined our approach using data from a well-studied diploid genome NA12878 that contains two alleles of each chromosome. We downloaded raw Illumina (300×) and Pacbio reads (31×) from the Genome In A Bottle (GIAB) consortium (Zook et al. 2014) and assessed results based on the NCBI build v38 assembly (**Methods**). In total, our approach identified 59,640 SVs, including 5,801 large deletions (> 50bp), 18,418 small deletions (≤ 50bp), 9,299 large insertions (> 50bp) and 26,122 small insertions (≤ 50bp) (**Supplemental Table S1**).

Pendleton et al. (Pendleton et al. 2015) recently performed *de novo* whole genome assembly of NA12878 using Celera Assembly (Myers et al. 2000) and Falcon from error-corrected Pacbio reads, followed by scaffolding with genome maps produced by Bionano technology and phasing with Illumina and Pacbio reads. This assembly can be used as a gold standard to assess the accuracy of SV assemblies.

### Large deletions in NA12878

For large deletions, we created a gold standard deletion set (referred below as GD) by merging five deletion call-sets produced respectively by 1) our HySA approach, 2) Delly (Rausch et al. 2012), 3) PBHoney (English et al. 2014), 4) a customized pipeline (CP) in Pendleton et al. (Pendleton et al. 2015) and 5) svclassify (Parikh et al. 2016), and validating each deletion at sequence resolution using the Pendleton et al. assembly (Pendleton et al. 2015) (**Methods**). The svclassify call-set was a high confidence set obtained from Personalis and the 1000 Genomes Project (Consortium 2010; Mills et al. 2011) and was the result of a machine learning method integrating signals in Illumina, Pacbio and Moleculo reads, and thus also resulted from a hybrid approach.

We plotted the receiver operating characteristics (ROC) curves based on the comparison of each of these 5 sets with the GD set (**Fig. 2**). By a fairly large margin, HySA outperformed Delly, PBhoney, CP and svclassify. The svclassify call-set is slightly inferior to HySA (specificity difference < 0.1 at a sensitivity of 0.33) even though it incorporated data from additional Moleculo long reads. In summary, our HySA approach achieved evidently better accuracy than approaches based on single technologies such as Delly, PBHoney and CP, and was favorable over another hybrid approach.

### Large insertions in NA12878

Among the 1,672 large (> 500bp) insertions we detected, 783 cannot be properly aligned to the NCBI build v38 assembly (**Methods**). Among them, 642 can be aligned to the Pendleton et al. assembly or the fosmid clones of NA12878 (Kidd et al. 2008). Only 141 (8.4%) had no supporting evidence from available data.

### Small INDELs in NA12878

We compared our small INDEL set (≤50 bp) with the Platinum (Eberle, M. A., E. Fritzilas, P. Krusche, M. Källberg, B. L. Moore, M. A. Bekritsky, Z. Iqbal, H.-Y. Chuang, S. J. Humphray, A. L. Halpert et al. 2016) and the GIAB set (**Fig. 3**). Interestingly, for the small insertions, we observed more overlapping calls between our set and the GIAB set than between the Platinum set and the GIAB set. In addition, we discovered 10,881 novel insertions that were neither in the Platinum nor in the GIAB sets. A majority of these novel insertions overlap with the dbSNP database (7,770) or the known STRs (6,835), indicating their potential validity.

**Fig. 3.**
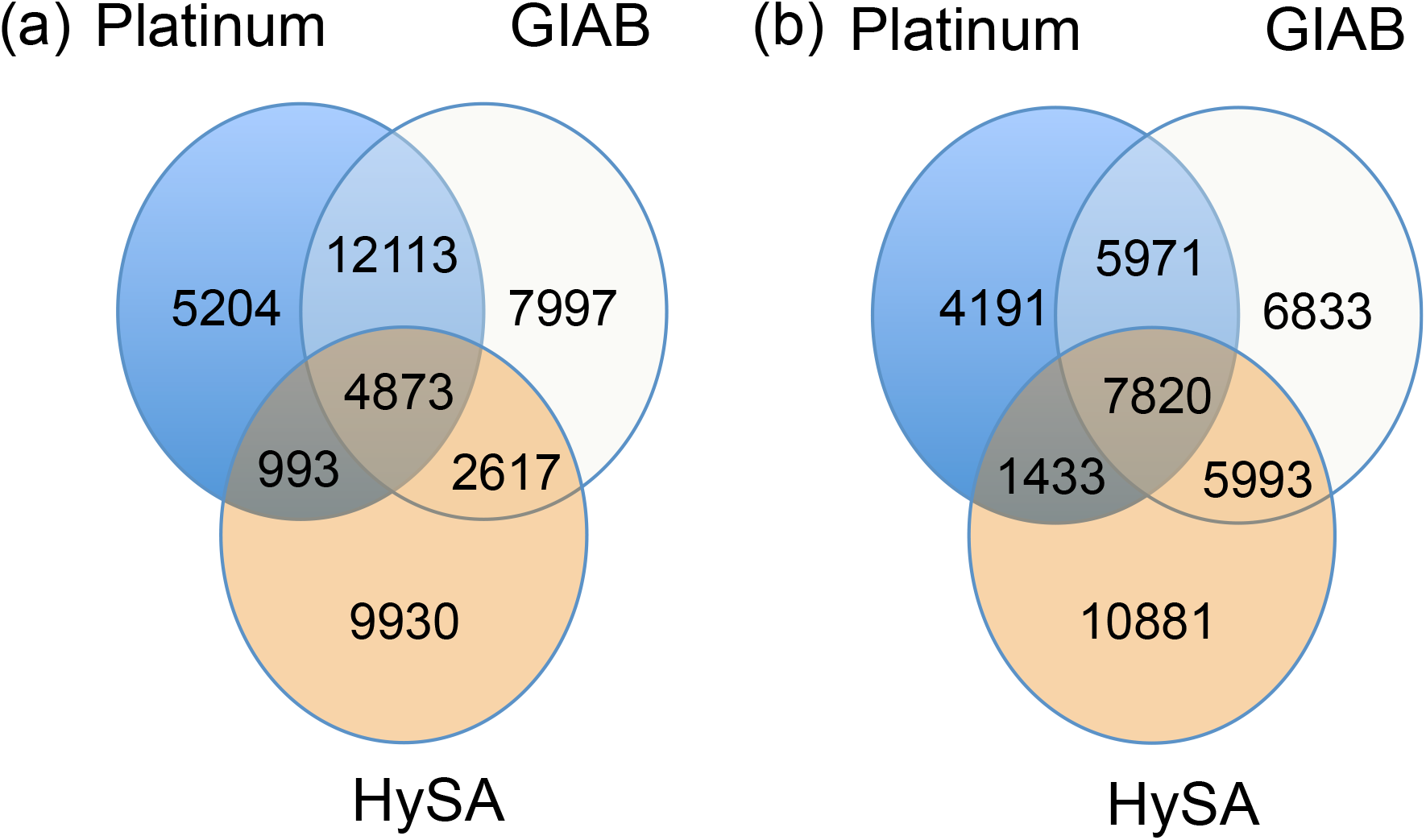
Comparison of small INDELs detected in NA12878 by HySA with those in the Platinum and GIAB sets. (a) Deletions with 1-bp overlap criterion. (b) Insertions with 1-bp overlap criterion after padding 50bp on the left and right of the insertion breakpoints.

For the small deletions, a large number in our set were shared with the Platinum and the GIAB sets. Of the 9,930 deletions that were unique to our set, 5,792 overlapped with dbSNP and 8,855 overlapped with the known STRs, indicating their potential validity. Manual inspection of our novel calls indicated that they were likely missed due to insufficient coverage or lack of alignment accuracy in a single source.

### Coverage Analysis

To provide a guideline for experimental design, we estimated the discovery sensitivity of our algorithm as a function of the Illumina coverage from 30× to 300× and the Pacbio coverage from 5× to 30× on chromosome 20 (**Fig. 4**, **Methods**). The sensitivity was defined as the fraction of the large deletions that our algorithm detected in the GD set. As a reference, we also computed the sensitivity of Delly from the Illumina data. As expected, the sensitivities increased with the increase of Pacbio or Illumina coverage. However, the gain of sensitivity decreased with the increase of Pacbio coverage, and the largest gain (∼0.25) was observed from 5× to 10× at all the inspected Illumina coverage.

**Fig. 4.**
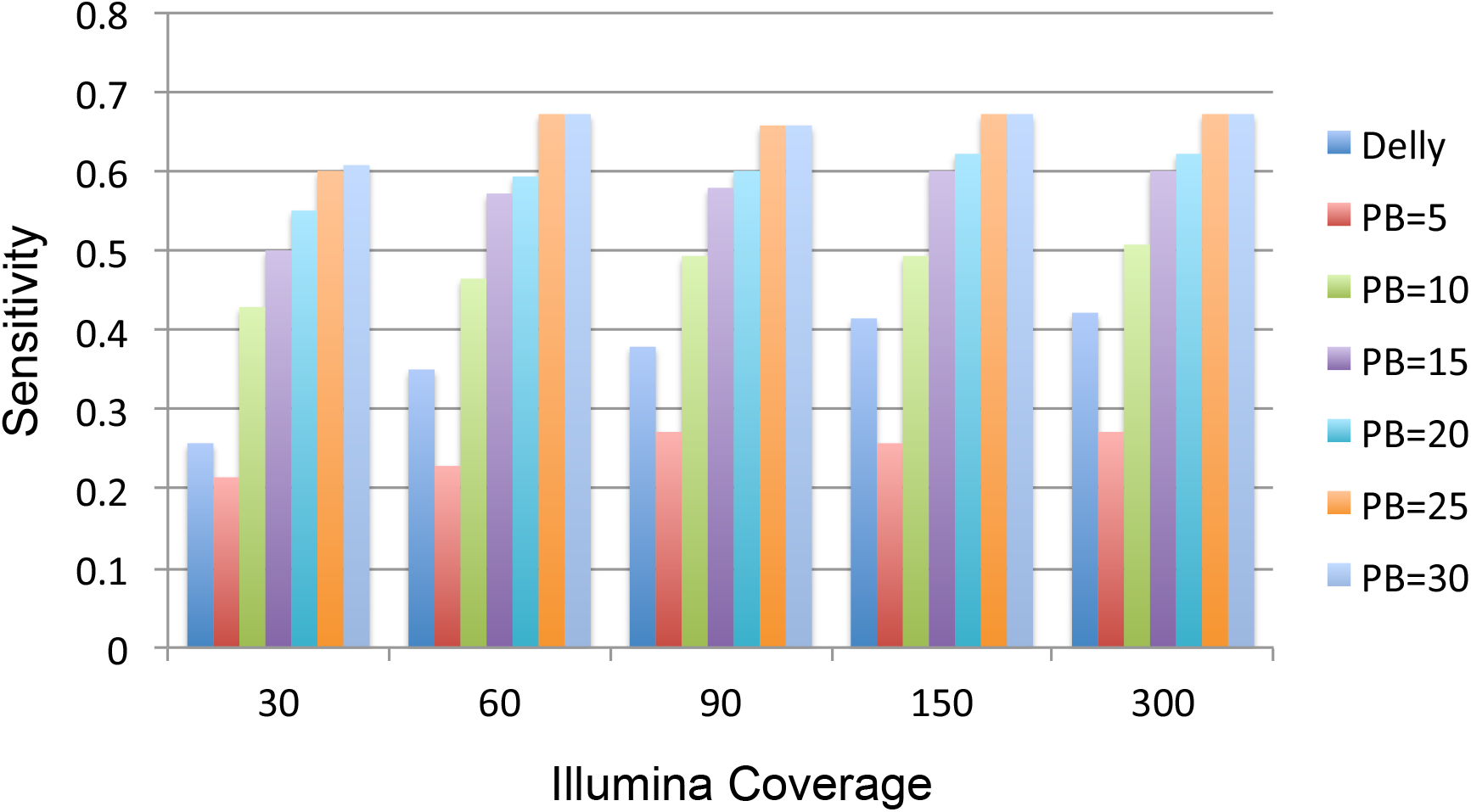
Coverage analysis. Sensitivities of our HySA approach are estimated at combinations of 5, 10, 15, 20, 25 and 30× Pacbio coverage and 30, 60, 90, 150 and 300× Illumina coverage, respectively. Sensitivities of Delly at given Illumina coverage are also shown on the leftmost bar.

Pacbio coverage no longer benefited the sensitivity to a great extent after it reached 25×. Likewise, the gain of sensitivity decreased with the increase of Illumina’s coverage, and saturation was observed at 60× when Pacbio’s coverage reached 25×. Therefore, a likely optimal combination of coverage would be 60× Illumina and 25× Pacbio reads, in order to obtain the most cost-effective hybrid SV assembly in a diploid genome using our approach.

In comparison, Delly has a higher sensitivity than our algorithm when Pacbio coverage is lower than 5× regardless of the Illumina coverage. However, once Pacbio coverage reaches 10× or above, our algorithm achieved higher sensitivity. This is expected because our algorithm requires at least 10× Pacbio coverage to successfully assemble the heterozygous deletions in a diploid genome. Notably, with 10× Pacbio coverage and 30× Illumina coverage, our algorithm achieved sensitivity comparable to Delly at 150× Illumina coverage. On the other hand, our algorithm achieved higher sensitivity at 10× Pacbio and 30× Illumina coverage than PBhoney at 30× Pacbio coverage. Given the current lower throughput and higher cost of Pacbio data, our approach clearly provides a more cost-effective solution than approaches that utilize only Illumina or Pacbio data but

## Discussion

In this work, we developed a new approach that performs targeted hybrid SV assembly from NGS and SMS reads for SV detection. Our approach combines the advantages of two technologies, the accuracy of the Illumina reads and the length of the Pacbio reads and was able to discover novel SVs missed by algorithms that detect SVs from a single technology, or by naively merging platform-specific call-sets. Our approach complements existing approaches and can be applied to substantially enhance the discovery power of ongoing personal genomic projects. The FDRs of our approach appeared to be low (<10%), owing partly to the combined use of orthogonal technologies. Although our approach was developed and assessed using data produced by Illumina and Pacbio technologies, the general framework is potentially applicable to data produced by other NGS and SMS technologies such as Ion Proton and Oxford Nanopore.

Dramatically different error profiles and lengths between sequencing reads generated by different technologies made it difficult to perform hybrid assembly using standard approaches such as OLC, *de Bruijn* and string graphs. The graph-theoretic approach that we developed and assessed in this study appears effective for constructing accurate hybrid SV assembly. In addition, focusing on SVs that are difficult to assemble by the WGA approaches results in computational efficiency and applicability of our algorithm in translational research.

We quantified the advantage of having both Illumina and Pacbio coverage in an assembly project and found that a combination of 25× Pacbio coverage and 60× Illumina coverage is likely optimal for comprehensively assembling a diploid genome and that at least 10× coverage of Pacbio reads is required for our HySA approach to perform well. This requirement appeared to result from a limitation of the Celera Assembler that we used to perform the local assembly. New tools under development may alleviate such limitations. For example, CANU (Berlin et al. 2015) as a fork of Celera Assembly has been developed to assemble high-noise single-molecule sequencing and is more capable of assembling lower coverage (< 10x) Pacbio data. Employing these new assemblers in HySA may further enhance the discovery of heterogeneous SVs, particularly those represented in low coverage (e.g., sub-clonal SVs in tumor bulk issue sequencing). Although the relationship we revealed between discovery power and coverage is specific to our algorithm, it is potentially generalizable to any discovery approaches that utilize both Illumina and Pacbio data.

Our work further highlights the complexity of human genomes and limitations of current technologies and approaches. Clearly, to obtain a perfect genome assembly and detect all the SVs, multiple technologies and computational algorithms that are advantageous in complementary ways will have to be employed synergistically. Despite the combined use of Pacbio and Illumina technologies, a substantial portion of SVs was likely undetected, particularly in highly repetitive areas of the genomes. Overcoming such limitation will require further development of sequencing technologies, as well as hybrid approaches that leverage the unique strengths of each technology.

## Methods

The two major parts of our algorithm, 1) clustering and 2) assembly/SV calling are described in Box 1 and 2.

Let ℐ be a set of Illumina reads and 𝒫 be a set of Pacbio reads. The overall objective is to identify for each unknown SV location *x*, the subsets *I* ⊂ ℐ and *P* ⊂ 𝒫 of Illumina and Pacbio reads that contain *x*, assemble the genomic region around *x* using *I* and *P*, and recover *x*. To achieve this objective, we propose a two-step solution in which the first step clusters the Illumina and Pacbio reads by SV sequences, and the second step conducts the assembly and SV calling from the clusters. A cluster is the pair (*I*,*P*) that corresponds to one potential SV, as discussed above.

### Box 1.

#### Algorithm of clustering Illumina and Pacbio reads

##### Algorithm 1: Cluster

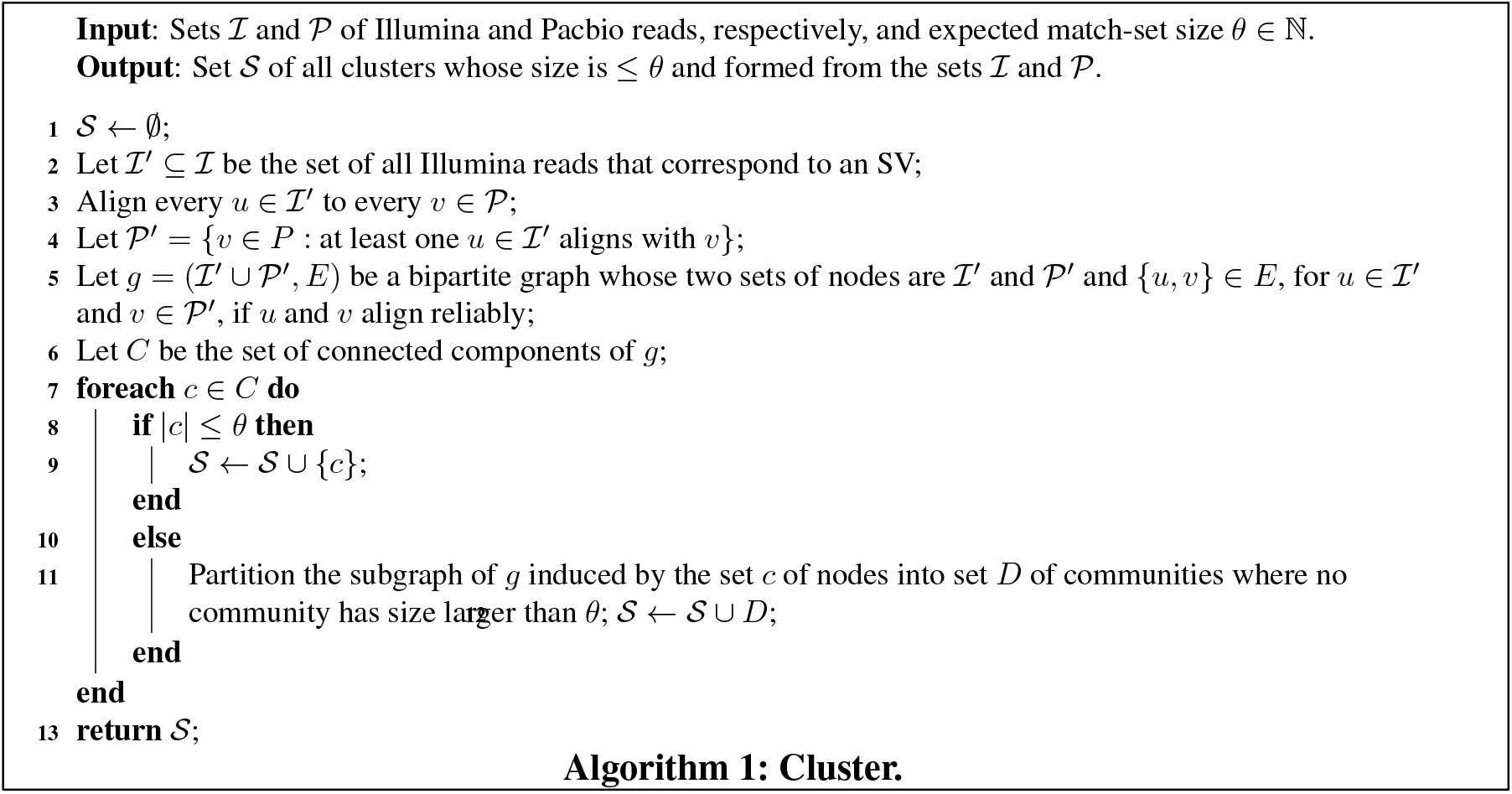

### Box 2.

#### Algorithm of inferring structural variation

##### Algorithm 2: ComputeSV

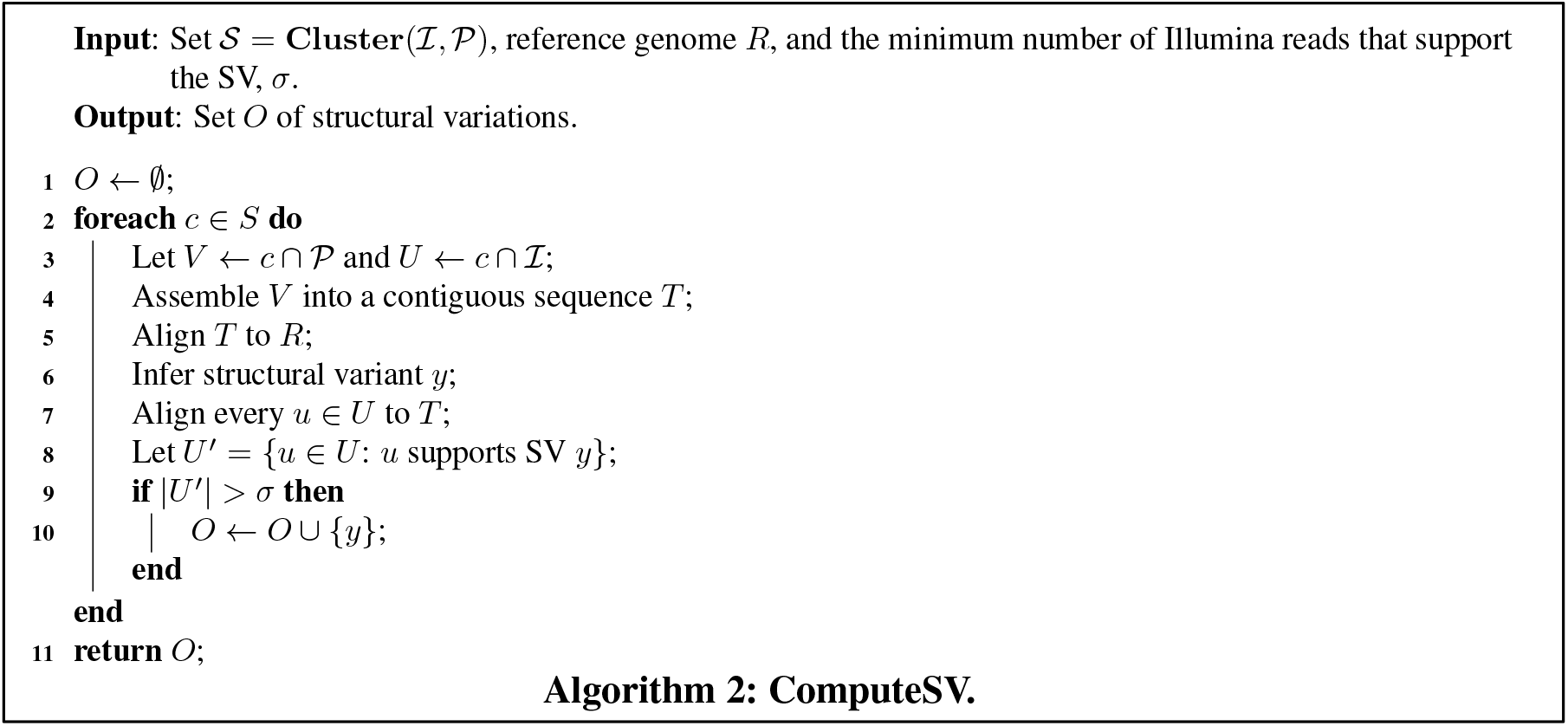

### Detailed description of steps in Algorithm 1

#### Step 2 (Extracting Illumina reads)

After aligning Illumina reads to the reference by BWA mem (Li 2013) (0.7.5a-r405), the read pairs that are discordant, unmapped, or have at least one read clipped or containing a large gap are extracted. Given a pre-computed average insert size *m* and standard deviation *δ* from a random 10K base pair (bp) region from the whole genome alignment, a read pair is deemed discordant if one of the following three happens:

1. The read pair’s insert size is > *m* + 4*δ*;
2. The paired reads’ orientation or position is abnormal;
3. The paired reads are aligned to two chromosomes.

Clipped reads should meet the quality requirement (median base quality of the clipped sequence > 35). Discordant reads, unmapped reads and reads with a large gap should have the median base quality > 35. All reads should have median base quality of both ends (20bp) > 35, and mapping quality > 0 for mapped reads.

#### Step 3 (Aligning extracted Illumina reads to Pacbio reads)

After removing the hairpin adaptor sequence from Pacbio polymerase read with pls2fasta (Chaisson and Tesler 2012), we partition the resulting sub-reads into multiple fasta files so that each file is approximately the size of a reference genome. With a customized version of BLASR that has one more option (-minInterval) allowing the recovery of more hits, we aligned all extracted Illumina reads to each sub-read file (-nCandidates 40 - minInterval 40 –maxScore 0 –minMatch 4 –maxMatch 13 -bestn 5). BLASR was developed to align Pacbio reads to the reference. Our usage (aligning Illumina to Pacbio reads) of it, although not typical, has been proven effective in terms of alignment accuracy as compared with other aligners such as BWA (**Supplemental Fig. 1**).

#### Step 5 (Selecting Reliable Alignments)

Due to Pacbio reads’ high error rate, false alignment from an Illumina read to a Pacbio read may occur. To reduce the number of false alignment, we required 70% of Illumina read sequence to be aligned to Pacbio reads and 70% of read were non-clipped. Moreover, Illumina reads’ paired signal is used for selecting reliable alignments. The criteria are summarized as below.

1. The percentage of identity (PI) of both ends’ alignment is > 70%;
2. One of the reads has at least 70bp aligned sequence;
3. The read pair is aligned concordantly with the same criteria described in Step 1 (**Fig. 1c**).

On each Pacbio read, we expect a set of Illumina reads aligned and piled to a certain location, where the breakpoint lies. Excessive Illumina reads piled together indicates a potential repetitive region. On the other hand, a small number of Illumina reads piled at one location indicates potential false alignment. We remove the alignments (**Fig. 1d**) involved in these two situations by setting up a range (3,*λK_I_*) in which *K_I_* is the mean coverage of Illumina reads, and *λ* is a threshold set by the user (heuristically, a reasonable *λ* is within (1, 1.2)).

In a diploid genome, Pacbio reads corresponding to the reference genome can be falsely extracted by clipped Illumina reads. To avoid extracting these Pacbio reads, we synthesize pseudo (a.k.a. “ref”) read pair from the reference sequence in identical positions and orientations whose sequence is a concatenation of the aligned subsequence of the Illumina read and the reference sequence corresponding to the clipped parts. The purpose of the “ref” read is to discern the allele the Pacbio read belongs to. Both clipped and pseudo (ref) reads are aligned to all Pacbio reads. A pseudo (ref) read is considered to align better if it spans longer (> 10bp) than its clipped counterpart, or its spanning sequence on the Pacbio read is close in length to its matched sequence (< 30bp) whereas the clipped read’s spanning sequence is not (**Fig. 1e**). When that happens, the alignment between the Illumina clipped read and the Pacbio read is regarded as false.

#### Step 6 (Computing the connected components of the bipartite graph)

The clustering algorithm Union-Find (Sedgewick and Wayne) is used to partition the bipartite graph into connected components (**Fig. 1b**). Due to the large distance of neighboring SVs on human genome (∼50,000bp/SV), the graph is sparse (∼6.5 million nodes and ∼11 million edges for a haploid genome CHM1 with 50× Illumina and 46× Pacbio), allowing Union-Find to have near-linear computational complexity.

#### Step 11 (Partitioning large connected components into communities)

Ideally one connected component corresponds to reads from one SV. Two factors may lead to reads from two or more SVs resulting in a single, large connected component: 1) false alignments due to sequence errors or presence of repetitive regions on the reference, and 2) multiple SVs in distances shorter than the read length. Such connected components are selected if their Pacbio read number is >*θ*, which is a pre-defined parameter as a function of Pacbio coverage. Due to the fact that Pacbio errors are random, and the homologous sequences differ from each other, though slightly, one potential solution for decomposing large connected components is to partition their induced subgraphs (in the bipartite graph) using a community detection algorithm (Newman 2006). Existing algorithms include Newman’s method (Newman 2006), Infomap (Rosvall and Bergstrom 2008), and mergeresplit (Zhou and Nakhleh 2012). Here we use Infomap (Rosvall and Bergstrom 2008) (version: 0.18.5) credited to its speed and ease-of-use (**Fig. 1b**).

### Detailed description of steps in Algorithm 2

#### Step 4 (Assembling Pacbio reads)

We use Celera Assembly (Myers et al. 2000) (8.3rc1) to assemble Pacbio reads in each cluster (ovlErrorRate=0.40 utgGraphErrorRate=0.40 cnsErrorRate=0.40 cgwErrorRate=040 unitigger=bogart obtErrorRate=0.30).

#### Step 5 (Aligning assembled contigs to the reference)

The assembled contigs are aligned to the reference with BLASR (Chaisson and Tesler 2012) (version: 1.3.1, parameter: -maxAnchorsPerPosition 100 –advanceExactMatches 10 –affineAlign –affineOpen 100 –affineExtend 0 –insertion 5 –deletion 5 –extend –maxExtendDropoff 20 –clipping subread –bestn 3).

#### Step 6 (Inferring structural variants)

The alignment of the contigs to the reference is ignored if either clipped end is > 500bp. The rest of the alignments are analyzed for search of large INDEL gaps (> 10bp). We require a matching flanking region > 10bp. For large gaps, BLASR tends to chop them into small ones separated by short matching subsequences. To accurately infer breakpoints for these gaps, we implemented a local re-alignment algorithm (pair-HMM (Durbin 1998)) from the assembled contig to the reference. The HMM has three states (‘M’ as Matching, ‘I’ as Insertion and ‘D’ as Deletion). Transitions are encouraged from ‘M’ to ‘M’, ‘D’ to ‘D’ and ‘I’ to ‘I’ with transition probabilities 0.99. Other transitions are discouraged with small transition probabilities (< 0.01). Through this process the small INDELs segmented by BLASR can be concatenated. We notice a similar procedure in MultiBreak-SV (Ritz et al. 2014) and Pendleton et al. (Pendleton et al. 2015).

#### Step 7 (Aligning Illumina reads in the same cluster to the assembled contig)

We align the Illumina reads in the same cluster to the contig that generates the SV using the customized BLASR (Chaisson and Tesler 2012) with parameters (-nCandidates 1 - minInterval 40 –maxScore 0 –minMatch 4 –maxMatch 13 -bestn 1). We pair up Illumina reads and filter mis-alignments by the same procedure as we described in Algorithm 1 Step 5.

#### Step 8 (Confirming structural variant)

In determining deletions, we require that the Illumina reads’ sequence matches 10bp on the left and right flanking region of the breakpoint with > 70 percentage of identity. In confirming insertion calls, we require that the Illumina reads’ inserted sequence matches that of Pacbio reads’ inserted sequence with > 70 percentage of identity.

### Reference alignment of Illumina reads for NA12878

BWA mem (0.7.5a-r405, with options -M -T0) was used to align the Illumina reads to NCBI build v38 assembly.

### BLASR reference alignment of Pacbio reads

We use the following options of BLASR for reference alignment of the CHM1 data (-bestn 24 -maxAnchorsPerPosition 100 -advanceExactMatches 10 -affineAlign - affineOpen 100 -affineExtend 0 -insertion 5 -deletion 5 -extend -maxExtendDropoff 20 - clipping subread -clipping soft -nproc 24).

### Simulation for evaluating the accuracy of the bipartite graph

The simulation tool wgsim (Li et al. 2009) was used to simulate Illumina paired end reads. We use PBSIM (Ono et al. 2013) (1.0.3, with options --difference-ratio 5:75:20 --length-mean 12000 --accuracy-mean 0.85 --model_qc model_qc_clr; model_qc_clr was provided by PBSIM package) to simulated Pacbio reads. The alignment from Illumina reads Pacbio reads was done by BWA mem (0.7.10 with option -x pbread) and BLASR (customized version, with options -minInterval 40 -nCandidates 40 -maxScore 0 - minMatch 4 -maxMatch 13. -bestn is twice of the Pacbio coverage) followed by filtering (> 87% percentage of identity, > 87% aligned sequence on one end).

For a pair of clusters (*μ,ν*), each containing a set of reads, *μ* from the inferred clustering,*v* from the ground truth, we computed the Jaccard index 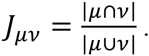 For each cluster *v* from the ground truth, we found its largest Jaccard index from all the clusters in the inferred clusters and assigned it to be *v*’s Jaccard index. We defined the 10% quantile of the Jaccard indices as JI90. By this definition, 90% of the clusters *v* have Jaccard indices greater than or equal to JI90.

### Generating curated large deletion calls (>50bp) for CHM1 and NA12878

In the CHM1 experiments, we ran Delly (0.7.2, default parameter) on the 50× Illumina data and combined them with those from Chaisson et al (Chaisson et al. 2015b) into one set. For each call in this set, we reconstructed the alternative allele by concatenating the sequences from two breakpoints on the reference (NCBI build v37). We extracted soft-clipped (≥10 clipped bases) and unmapped Illumina reads that have positive mapping quality and <3 mismatches falling within [-500, 50] bp of the left breakpoint and [-50, 500] bp of the right breakpoint to the alternative allele and aligned (BWA mem 0.7.5a-r405, default parameter) them to the reconstructed alternative alleles. If an Illumina read aligns to the reconstructed alternative allele without soft clipping and simultaneously spans the breakpoint, it is counted as genotyping the call. A call is curated if it is genotyped by at least two Illumina reads.

In the NA12878 experiments, we downloaded calls generated by svclassify, PBHoney and a customized pipeline. We ran Delly (0.7.2, default parameter) on the 300X Illumina bam. The five call sets including ours were combined and de-duplicated. For each call in this union set, we reconstructed the alternative allele by concatenating sequences from two breakpoints on the reference (NCBI build v38) in the same manner as that in CHM1. The reconstructed alternative allele was aligned (BWA mem 0.7.5a-r405, default parameter) to the NA12878 whole genome hybrid assembly (Pendleton et al. 2015). A call is counted as validated if the alignment meets the same criterion as that in validation of CHM1 deletion calls by CHM1_1.1.

### Calculating large deletion signals for CHM1

In calculating variant-supporting Illumina read number, we used the same strategy as the one described for generating curated calls in CHM1 except that we extracted all soft clipped and unmapped reads and align them to the reconstructed alternative allele. In calculating variant-supporting Pacbio read number, for each call, we selected Pacbio reads overlapping with [-100, 100] bp of the left breakpoint whose alignment to the reference had the total length of the clipped end 1) < 500bp, and 2) < 0.2 of the total read length. A read is counted as supporting the variant if it has a deletion gap starting within 300bp of the inferred left breakpoint, and the deletion size is within 30bp of the inferred SV size.

### Estimating FDR for Large Deletion Calls by CHM1_1.1

To evaluate the false discovery rate of our large deletion calls, we reconstructed the alternative allele from our calls by concatenating the sequences (500bp on each end) from two breakpoints on the reference (NCBI build v37) and aligned (BWA mem, 0.7.5a-r405, default parameter) the concatenated sequence (total length = 1000bp) to the Berlin et al. whole genome assembly (Berlin et al. 2015). The calls that had both the largest gap and clipped sequence < 50bp, and the total length of all gaps and clipped sequence < 100bp was counted as validated. Among the calls that cannot be directly validated by the Berlin et al.assembly, 63/450 overlapped with the curated call set. For the rest of the 387 calls, we extracted soft-clipped Illumina reads from [-500, 50] of the left breakpoint and [-50, 500] of the right breakpoint and aligned them to the alternative allele. There were 160 calls that had >= 2 Illumina reads that were not clipped and spanned the breakpoint. Treating these two sets of calls as validated, a likely more accurate estimate of the FDR is < 7.5% (227/3007).

### Validating insertion for CHM1 and NA12878

For each insertion call, we extracted inserted sequence and aligned it to the reference (BWA mem, 0.7.5a-r405). For CHM1, the reference includes NCBI build v37, build v38, Berlin et al. whole genome assembly (Berlin et al. 2015) and assembled contigs from Chaisson et al. Duplicate calls with the same contig were not included in the calculation. The 29 insertions calls validated by build v38 or the whole genome assembly but not by Chaisson et al. were manually inspected by samtools tview (Li et al. 2009) of the Illumina reads. For NA12878, the reference includes NCBI build v38, the hybrid assembly (Pendleton et al. 2015) and fosmid clones of NA12878 (Kidd et al. 2008). For both CHM1 and NA12878, if an insertion has an alignment with the largest gap < 50 bp, clipped sequence length < 100 bp, and the matched sequence > 0.9 of the total sequence length, it is counted as validated.

### Comparison with Pindel and GATK

Pindel (0.2.5) was run with default parameter with a post filtering of ≥5 supporting split reads. GATK was run on three steps: HaplotypeCaller (--genotyping_mode DISCOVERY -stand_emit_conf 10 -stand_call_conf 30), SelectVariants (-selectType INDEL) and VariantFiltration (--filterExpression "QD < 2.0 ║║ FS > 200.0 ║║ ReadPosRankSum < -20.0" --filterName "indel_filter").

### Comparison criteria on call sets

In both CHM1 and NA12878 analysis, 50% reciprocal criterion was used to overlap large deletions. The 1-bp overlapping criterion was used in comparing small INDELs. For all insertions, 50bp were padded on the left and right of the inserted breakpoints to account for ambiguity in breakpoint locations in repeats.

### Coverage Analysis

We downsampled extracted Illumina reads to 150×, 90×, 60×, 30×, and Pacbio reads to 25×, 20×, 15×, 10×, 5×. For each coverage combination, the bipartite graph was built and partitioned into small clusters. For each validated call, if at least 5 Pacbio reads remained in a cluster corresponding to that from the highest coverage (300× for Illumina and 31× for Pacbio), it is counted as a hit. For comparison, we downsampled all Illumina reads that aligned to chromosome 20 to 150×, 90×, 60× and 30×, and ran Delly (0.7.2, default parameter) on the bams generated at each coverage.

### Software availability

The developed pipeline and the scripts used in this manuscript are online at https://bitbucket.org/xianfan/hybridassemblysv/overview.

### Data

CHM1 raw Pacbio data were downloaded from https://s3.amazonaws.com/datasets.pacb.com/2014/Human54x/raw/human54x_set[0-34].tgz.

CHM1 Illumina reads were downloaded from NCBI GenBank with accession number SRX652547.

CHM1 Chaisson et al. large deletion calls and large insertion sequences were downloaded from http://eichlerlab.gs.washington.edu/publications/chm1-structural-variation/.

CHM1_1.1 was downloaded from ftp://ftp.ncbi.nlm.nih.gov/genomes/all/GCA_000306695.2_CHM1_1.1/GCA_000306695.2_CHM1_1.1_genomic.fna.gz

NA12878 Pacbio data was downloaded from NCBI GenBank with accession numbers SRX627421 and SRX638310.

NA12878 Illumina data was downloaded from ftp://ftp-trace.ncbi.nlm.nih.gov/giab/ftp/data/NA12878/NIST_NA12878_HG001_HiSeq_300x/

NA12878 svclassify data was downloaded from ftp://ftp-trace.ncbi.nlm.nih.gov/giab/ftp/technical/svclassify_Manuscript/Supplementary_Information/Personalis_1000_Genomes_deduplicated_deletions.bed.

NA12878 sequences from fosmid clone were downloaded from GenBank Bioproject 29893, selected with ID ABC12.

NA12878 Platinum INDEL calls were downloaded from ftp://platgene_ro@ussd-ftp.illumina.com/hg19/8.0.1/NA12878/NA12878.vcf.gz

NA12878 GIAB INDEL calls were downloaded from ftp://ftp-trace.ncbi.nih.gov/giab/ftp/release/NA12878_HG001/NISTv3.2.2/NA12878_GIAB_highconf_IllFB-IllGATKHC-CG-Ion-Solid_ALLCHROM_v3.2.2_highconf.vcf.gz

NA12878 hybrid assembly was downloaded from ftp://ftp.ncbi.nlm.nih.gov/genomes/all/GCA_001013985.1_ASM101398v1/GCA_001013985.1_ASM101398v1_genomic.fna.gz

## Authors' contributions

XF, KC conceived the study. XF, MC designed and implemented the code. XF performed the analysis. XF, LN, and KC wrote the manuscript. KC and LN provided oversight and coordinated the project. All authors read, revised and approved the final manuscript.

## Acknowledgements

This work was supported in part by the National Cancer Institute (NCI) grant R01-CA172652 to K.C., National Human Genome Research Institute (NHGRI) grant U41-HG007497-01, and the National Cancer Institute Cancer Center Support Grant P30-CA016672. We would like to thank E. Eichler, J. Korbel, C. Lee and other members of the Human Genome Structural Variation Consortium for ideas and discussions. We would also like to thank W. Zhou for suggesting community decomposition algorithms, as well as Z. Chong for discussion of ideas and the update of BWA as well as graphic algorithms.

